# Assessing the effect of insecticide-treated cattle on tsetse abundance and trypanosome transmission at the wildlife-livestock interface in Serengeti, Tanzania

**DOI:** 10.1101/2020.04.14.040873

**Authors:** Jennifer S. Lord, Rachel S. Lea, Fiona K. Allan, Mechtilda Byamungu, David R. Hall, Jessica Lingley, Furaha Mramba, Edith Paxton, Glyn A. Vale, John W. Hargrove, Liam J. Morrison, Stephen J. Torr, Harriet K. Auty

## Abstract

In the absence of national control programmes against Rhodesian human African trypanosomiasis, farmer-led treatment of cattle with pyrethroid-based insecticides may be an effective strategy for foci at the edges of wildlife areas, but there is limited evidence to support this.

We combined data on insecticide use by farmers, tsetse abundance and trypanosome prevalence with mathematical models to quantify the likely impact of insecticide-treated cattle.

Sixteen percent of farmers reported treating cattle with a pyrethroid, and chemical analysis indicated 18% of individual cattle had been treated, in the previous week. Treatment of cattle was estimated to increase daily mortality of tsetse by 5 – 14%. Trypanosome prevalence in tsetse, predominantly from wildlife areas, was 1.25% for *T. brucei s.l*. and 0.03% for *T. b. rhodesiense*. For 750 cattle sampled from 48 herds, 2.3% were PCR positive for *T. brucei s.l.* and none for *T. b. rhodesiense*. Using mathematical models, we estimated there was 8 – 29% increase in mortality of tsetse in farming areas and this increase can explain the relatively low prevalence of *T. brucei s.l.* in cattle.

Farmer-led treatment of cattle with pyrethroids is likely, in part, to be limiting the spill-over of human-infective trypanosomes from wildlife areas.

**Author summary:** The acute form of sleeping sickness in Africa is caused by the parasite *Trypanosoma brucei rhodesiense*. It is transmitted by tsetse flies and can be maintained in cycles involving both livestock and wildlife as hosts. Humans are incidentally infected and are particularly at risk of infection near protected areas where there are both wildlife and suitable habitat for tsetse. In these regions, the tsetse vector cannot be eradicated, nor can infection be prevented in wildlife. Here we use field studies of tsetse and livestock in combination with mathematical models of tsetse population change and trypanosome transmission to show that use of pyrethroid-based insecticides on cattle by farmers at the edge of protected areas could be contributing to lowering the risk of sleeping sickness in Serengeti District, Tanzania. To our knowledge, our study is the first to report farmer-led tsetse control, co-incident with tsetse decline and relatively low prevalence of *T. brucei s.l.* in cattle.

## Introduction

In East and Southern Africa, tsetse flies (*Glossina* spp) transmit *Trypanosoma brucei rhodesiense*, which causes Rhodesian human African trypanosomiasis (r-HAT). Tsetse also transmit *T. congolense, T. vivax* and *T. brucei*, the causative agents of animal African trypanosomiasis (AAT) in livestock.

*Trypanosoma brucei s.l., T. congolense* and *T. vivax* can circulate in transmission cycles involving livestock or wild mammals [1]. The extensive conservation areas of East and Southern Africa that support tsetse, as well as wildlife, can therefore be foci for r-HAT and AAT. At the interface of wildlife- and livestock areas, there is potential for trypanosomes to shift from a wildlife- to a livestock-dominated cycle of transmission [1]. Although existing r-HAT foci are often associated with wildlife areas, the importance of cattle as reservoirs at the wildlife-livestock interface is unclear [1].

There are few studies that address the role of cattle in r-HAT transmission in wildlife-livestock interface areas. Kaare et. al. [2] suggested that r-HAT could be re-emerging in Serengeti District, Tanzania, based on surveys of cattle adjacent to the Serengeti National Park in 2001, where they found 5.6% of cattle positive for *T. brucei s.l.* DNA and ~1% of 518 cattle sampled as positive for *T. b. rhodesiense* DNA.

With c. 1.4 million people living at moderate to high risk of *T. b. rhodesiense* in East and Southern Africa [3], there is a need to identify appropriate control measures that can reduce the risk of trypanosomiasis for both people and cattle living near wildlife areas. Previous modelling has indicated that insecticide-treated cattle could offer an effective method of control, particularly for r-HAT [4], but modelling has not been extended to consider wildlife-livestock interface areas.

We previously found that numbers of tsetse caught in traps declined by >90% across a wildlife-livestock interface in Serengeti District, Tanzania, with no tsetse being caught >5 km into farming areas [5]. Our previous work showed that this was due, in part, to reduced availability of habitat suitable for tsetse. This is likely to be typical for other r-HAT foci in and near wildlife areas, where increasing human and livestock densities lead to a reduction in tsetse habitat. However, the effect of habitat did not fully explain the change in tsetse abundance [5]. At the same time, we obtained preliminary evidence that livestock farmers were frequently treating their cattle with pyrethroids, insecticides effective against tsetse [6]. It seems likely that mass treatment of cattle with insecticide is reducing the density of tsetse populations and hence trypanosomes.

We aimed to assess whether the presence of insecticide-treated cattle is contributing to the decline in tsetse and quantify the impact of such a decline in tsetse on the transmission of trypanosomes in cattle at the interface between wildlife and livestock populations.

## Methods

### Study site

Our study site comprised the Serengeti National Park, adjacent game reserves and farming areas (S1 Fig). Farming areas are used predominately for livestock grazing and crop production, with c. 30 cattle/km^2^ [7].

The study site supports three species of tsetse – *G. swynnertoni*, *G. pallidipes* and *G. brevipalpis* [5]. The Serengeti area is an historic r-HAT focus [8]. Since the last outbreak in 2000/2001, during which at least 20 cases were reported in local populations and tourists [9,10], sporadic cases continue to occur [3].

### Tsetse surveys

We carried out surveys during February, June-July and October 2015 along four transects from 5 km inside wildlife areas, to 10 km into farming areas (S1 Fig). We set a total of 72 odour-baited Nzi traps, 38 inside wildlife areas and 34 outside, during each survey and emptied traps each day for three consecutive days, recording the sex and species of tsetse. Full details of the survey method are provided in Lord et al. (2018) [5].

We caught fewer than 100 *G. brevipalpis* during the study, so our analyses focussed only on *G. pallidipes* and *G. swynnertoni*. Since daily numbers (*y*) of tsetse caught per day in traps were overdispersed, we transformed the data to log_10_(*y* + 1) before calculation of average counts per trap.

During 2016 we carried out additional trapping inside wildlife areas, up to 10 km from the boundary, to catch sufficient numbers of tsetse to quantify the prevalence of *T. congolense* savanna and *T. brucei s. l.* in tsetse. *T. congolense* presence was used as a proxy for AAT, being more prevalent than *T. vivax* in the study area [2].

During each survey in 2015 and 2016, we transported tsetse flies, preserved in ethanol in individual tubes, to the Liverpool School of Tropical Medicine and then processed them for the detection of trypanosome DNA (S1 Text).

### Livestock surveys

We carried out a cross-sectional livestock survey, in villages <5 km from the wildlife boundary, during July-August 2016. A total of 48 herds and 750 cattle were selected using a stratified selection method (S1 text). For each sampled animal, we collected blood from the jugular vein into PAXGENE tubes, and recorded details of age, sex and any treatments given in the last six months. We administered a questionnaire to each livestock keeper to collect information on current vector control practices. Questions included the date the animals were last treated with insecticide and the method of application. Blood samples were tested by PCR for the presence of *T. brucei* and *T. congolense* DNA (S1 Text).

In addition to asking farmers about use of insecticides, we also analysed hair for the presence of pyrethroids. Using disposable razors, we collected hair samples (0.04 g/animal) from the flank of four randomly-selected cattle within each herd, giving a total of 176 samples, which were sealed individually in foil bags. Cypermethrin and alpha-cypermethrin was extracted from each sample in acetone and assayed by gas chromatography-mass spectrometry (GC-MS) (S1 Text). This method can detect the presence of insecticide at 7 days post application, but not at 14 days [11].

### Ethics statement

Cattle sampling involved venous blood sampling and collection of hair samples (procedures classified as ’mild’ under UK Home Office regulations). Discussions regarding the veterinary sampling were undertaken with key administrative and community leaders to inform communities of the overall study and mobilise households to participate. Animals were sampled by veterinarians or trained paraveterinary workers. Jugular blood samples (10ml) were taken into sterile vacutainers and hair samples collected using a safety razor. The animals were restrained appropriately to minimise the time and distress involved in the process of sample collection. All sampling was undertaken under the supervision of a veterinarian. Ethical approval for this work was obtained from the SRUC Animal Experiments Committee and the Commission for Science and Technology (Costech) in Tanzania (permit number 2016-33-NA-2014-233).

### Data summary

We calculated the prevalence, and exact binomial 95% confidence intervals, for *T. brucei s. l., T. brucei rhodesiense* and *T. congolense* in cattle and tsetse as the percentage of individuals testing PCR positive for each trypanosome species and subspecies. For tsetse, this prevalence includes infected flies that might not be infectious.

To estimate the possible range of tsetse daily mortality attributable to insecticide-treated cattle, we assumed that any given tsetse fly contacts a vertebrate host either every two or every three days [12]. We then estimated the proportion of cattle treated, using information from hair sample analysis and questionnaire responses. We divided this proportion by the duration of the feeding cycle, assuming that a fly would die from contacting any host testing positive for insecticide [6]. Under the hypothesis that cattle were treated with insecticide, we could not estimate the proportion of bloodmeals from cattle – because, by assumption, those that had fed on treated cattle would not be caught for analysis. We therefore made the assumption that cattle were the only source of bloodmeals in farming areas [13].

### Modelling tsetse population dynamics across the wildlife-livestock interface

To estimate the additional tsetse mortality in farming areas, we developed a spatially-explicit model of tsetse population dynamics and fitted the model to the tsetse catch data.

We describe changes in numbers of pupae (*P*) and adult tsetse (*A*) in space and time using two recursion equations on a lattice (S2 Text). Parameters used are described in Table 1.

**Table 1.**
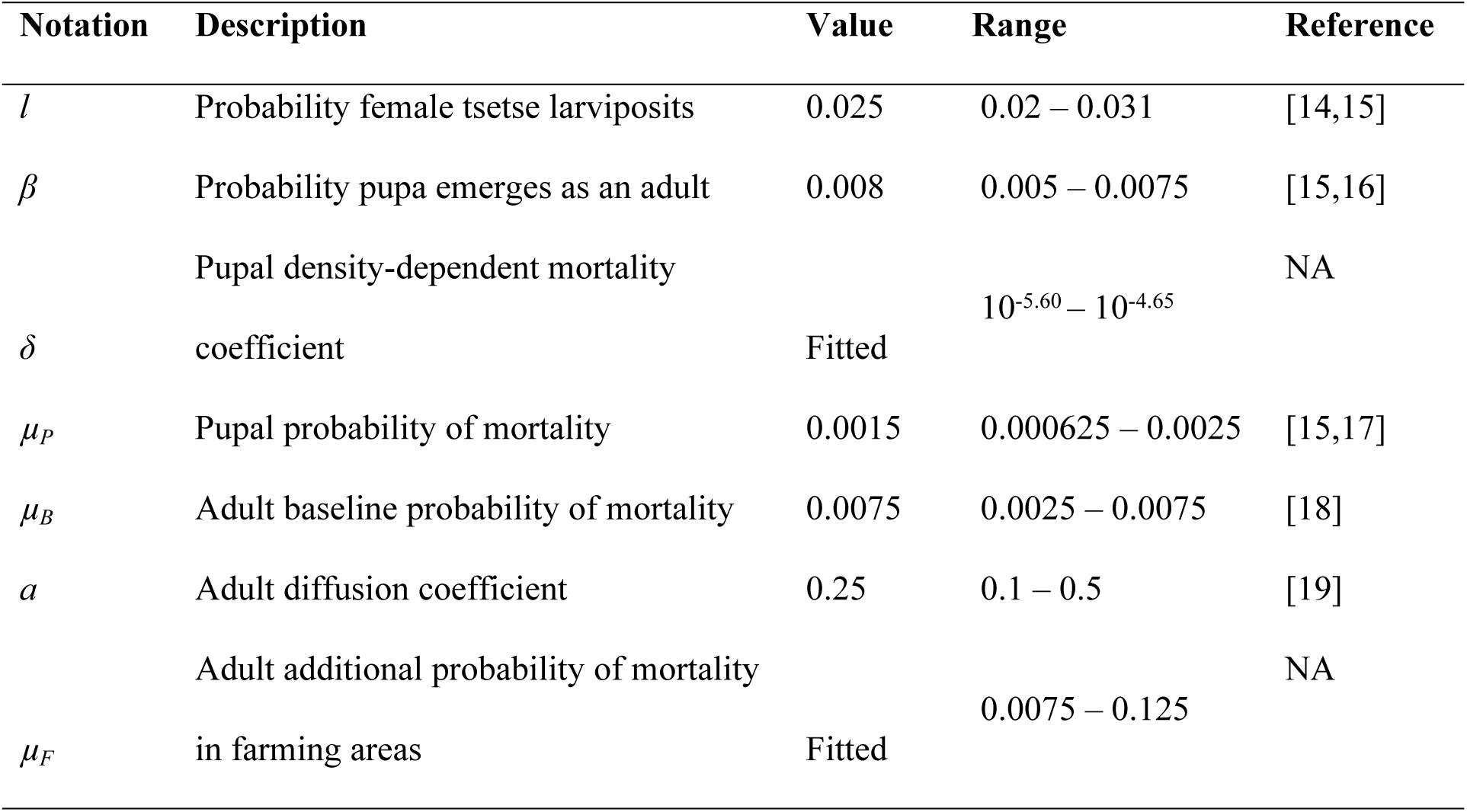
Parameters and values used in the model of tsetse population dynamics. Values are per 0.25 days. Each cell in the area modelled is a square of side 500 m.

Reflecting boundaries were used in the lattice so that for cells at the edge of the lattice, numbers of tsetse moving in were equivalent to those leaving. Each day, in each cell *i,j* a proportion *a* of adult tsetse diffuse into adjacent cells. Adult females, assumed to be half the population, produce pupae with probability *l*. Adults die with probability *μ*_*B*_. Pupae emerge as adults with probability *β* and are subject to density-independent (*μ*_*P*_) and density-dependent (*Pδ*) deaths. In addition to the baseline mortality, adults present in cells designated as ‘farming’ areas are subject to an additional mortality (*μ*_*F*_) to represent insecticide use and habitat degradation.

We carried out a sensitivity analysis (S3 Text), to quantify how the modelled decline in tsetse density across the wildlife-livestock interface was influenced by model parameter values. We then fitted the model to observed tsetse abundance data using nonlinear least squares regression implemented with the Levenberg-Marquardt Algorithm, accounting for uncertainty in parameter values (S3 Text).

### Modelling trypanosome transmission dynamics across the wildlife-livestock interface

To quantify the effect of tsetse population decline on trypanosome prevalence in cattle in the interface area, we extended the tsetse model to include trypanosome transmission (S2 Text).

In addition to tsetse population dynamics described above, adult tsetse in each cell progress through susceptible teneral (juvenile unfed) (*S*_*V*_) to either susceptible non-teneral (*G*_*V*_), or exposed (*E*_*1V*_ – *E*_*3V*_) and then infectious (*I*_*V*_) classes. Instead of having a fixed-time for the tsetse incubation period, or assuming that the incubation period is exponentially distributed, we model three exposed classes as per [14], assuming an Erlang distributed waiting time for the extrinsic incubation period [15]. Hosts in each cell progress through susceptible (*S*_*H*_), exposed (*E*_*H*_), infected/ infectious (*I*_*H*_) and recovered (*R*_*H*_) classes. We assumed that host populations do not move, and host birth and death rates are equal.

Due to uncertainty in parameter values (Table 2) for trypanosome transmission, to quantify the potential effect of the tsetse population decline on transmission across the interface, we first ran a sensitivity analysis without increased tsetse mortality (S3 Text). To determine the potential effect of increased tsetse mortality in farming areas on cattle trypanosome prevalence we selected from the sensitivity analysis combinations of parameter values that produced tsetse prevalence at equilibrium within the range observed in our study site for *T. brucei* and *T. congolense*. We then ran the model using the selected parameter combinations and including an additional tsetse mortality, the value of which we obtained from fitting the model of tsetse population dynamics to observed tsetse abundance.

**Table 2.**
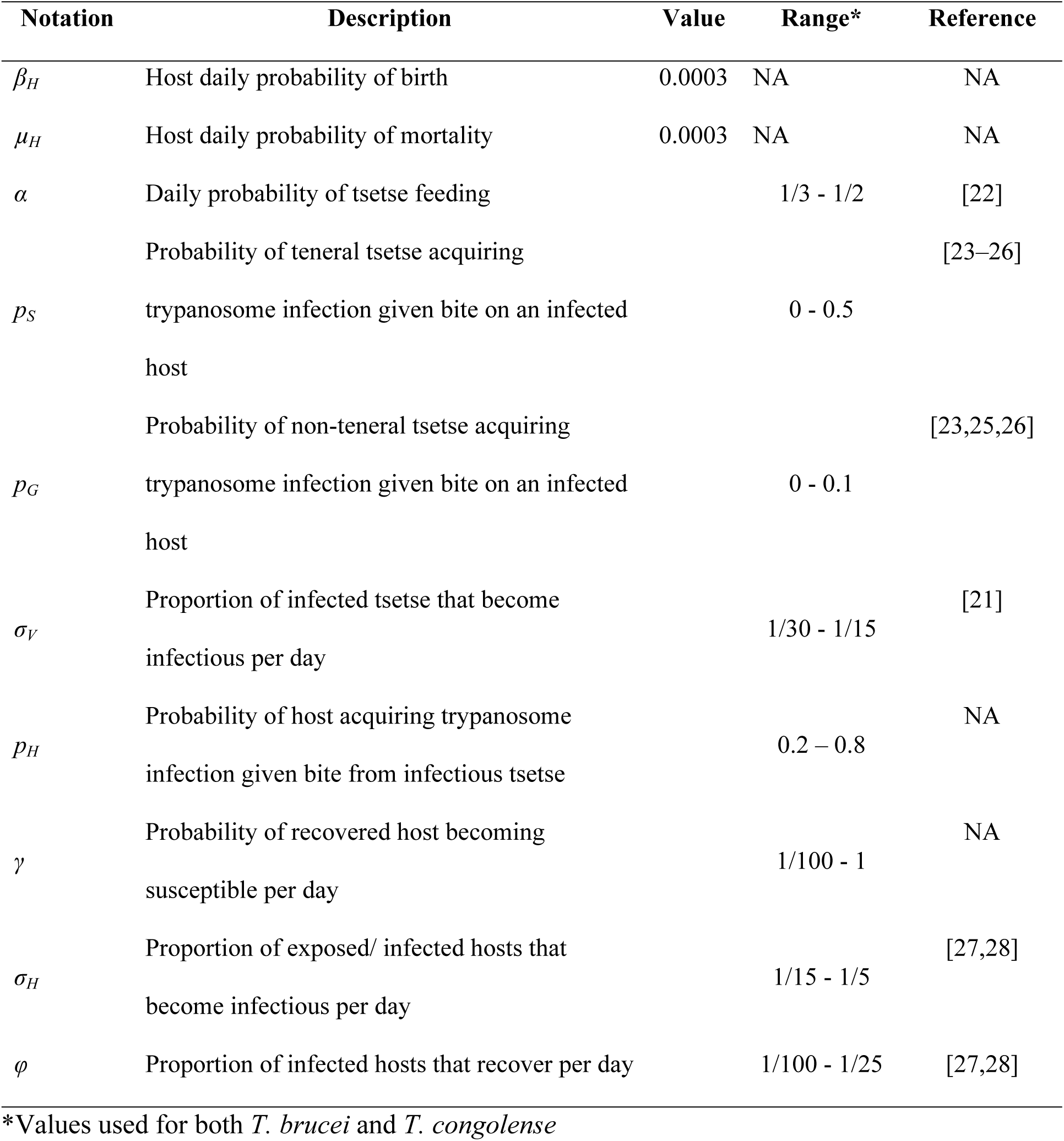
Parameters and values used in the trypanosome transmission model. See Table 1 for tsetse population dynamics parameters.

Both the tsetse population dynamics and trypanosome transmission models, plus code to produce the figures in this manuscript can be accessed at https://github.com/jenniesuz/tsetse_wli.git.

## Results

### Observed tsetse decline across the wildlife-livestock interface

Mean daily numbers of both *G. pallidipes* and *G. swynnertoni* caught per trap declined to zero by 5 km outside wildlife areas in the second and third quarterly surveys of 2015, similar to that observed during the first survey in February 2015 (Fig 1, [5]). Across all three surveys in wildlife areas, >99% of traps caught at least one tsetse, whereas in farming areas 58% of traps did not catch any flies.

**Fig 1.**
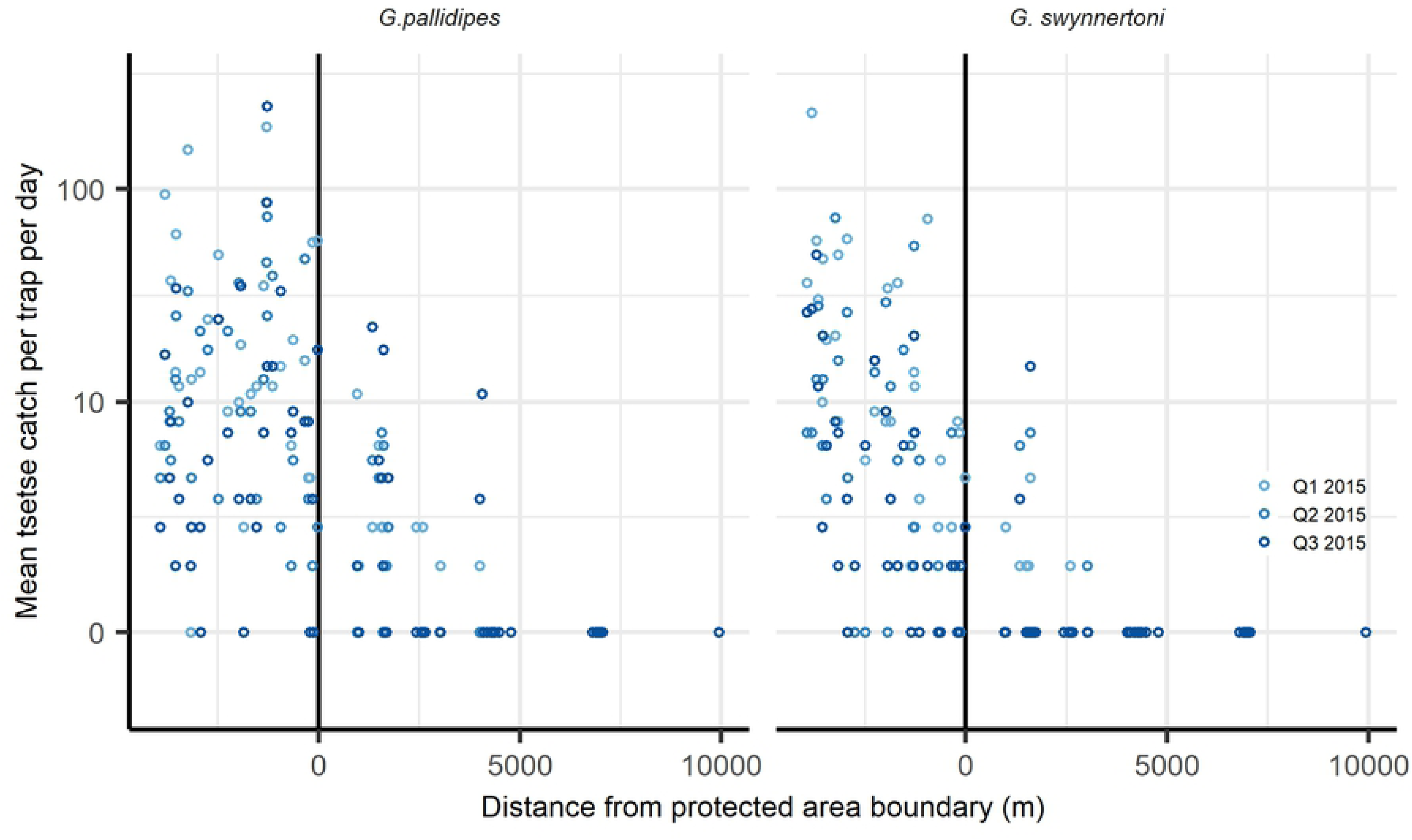
Mean numbers of tsetse caught across the wildlife-livestock interface by season and species during 2015.

### Observed trypanosome prevalence in tsetse and cattle

During 2015 and 2016 we caught 5986 tsetse, which were tested for the presence of trypanosome DNA. Only 4% flies sampled during 2015 were from farming areas. Both *T. congolense* and *T. brucei s.l.* were detected and two flies from wildlife areas tested positive for *T. b. rhodesiense* (Table 3). Of the 750 cattle sampled in 2016, none was positive for *T. b. rhodesiense* DNA and *T. brucei s.l.* prevalence was one seventh of that for *T. congolense* (Table 3).

**Table 3.** Prevalence of trypanosome species in tsetse and cattle. Prevalence defined as the percentage of hosts or vectors testing positive for the presence of DNA for the respective species: 95% confidence intervals in parentheses.

|  |  | Prevalence (%) |  |  |
| --- | --- | --- | --- | --- |
|  | No. sampled | *T. brucei rhodesiense* | *T. brucei s.l.* | *T. congolense* |
| **Tsetse** | 5986 | 0.03 (0.004 - 0.121) | 1.25 (0.09 - 1.57) | 5.34 (4.79 - 5.94) |
| **Cattle** | 750 | 0 (0 – 0.005) | 2.3 (1.3 - 3.6) | 16.7 (14.1 - 19.5) |

### Insecticide use

Of the 44 livestock owners questioned about insecticide use, 67% reported treating at least some of their cattle with a pyrethroid within the previous month and 16% reported treating within the previous week. Chemical analyses of hair samples collected at the time of the questionnaire showed that 18% of 176 individual cattle and 27% of 44 herds had detectable levels of alphacypermethrin or cypermethrin, indicating treatment within c. 7 days.

If we assume a three-day feeding cycle, and that 16% of cattle are treated weekly, tsetse mortality from insecticide-treated cattle would be c. 0.05 per day. If we assume a two-day feeding cycle and that 27% cattle are treated, mortality from insecticide would be c. 0.14 per day.

### Simulating tsetse population dynamics across the wildlife-livestock interface

We fitted the tsetse population dynamics model to mean tsetse catches per trap per day across all seasons, given that catches of both *G. pallidipes* and *G. swynnertoni*, across all seasons, declined to zero by 5 km outside wildlife areas (Fig 1).

Using the parameter values in Table 1, the best fit additional daily probability of adult mortality (*μ*_*F*_) was 0.15 per day (S1 Table, Fig 2). Of the fixed parameters, daily dispersal distance (*a*) and daily probability of larviposition (*l*) had the biggest influence on the relative density of tsetse 1 km inside farming areas, compared to density 5 km inside wildlife areas, with PRCC > 0.5 and < −0.5, respectively (S2 Fig, S3 Fig). Depending on values for the daily probability of larviposition and dispersal, fitted values for additional daily probability of mortality varied between 0.08 and 0.29 (S1 Table).

**Fig 2.**
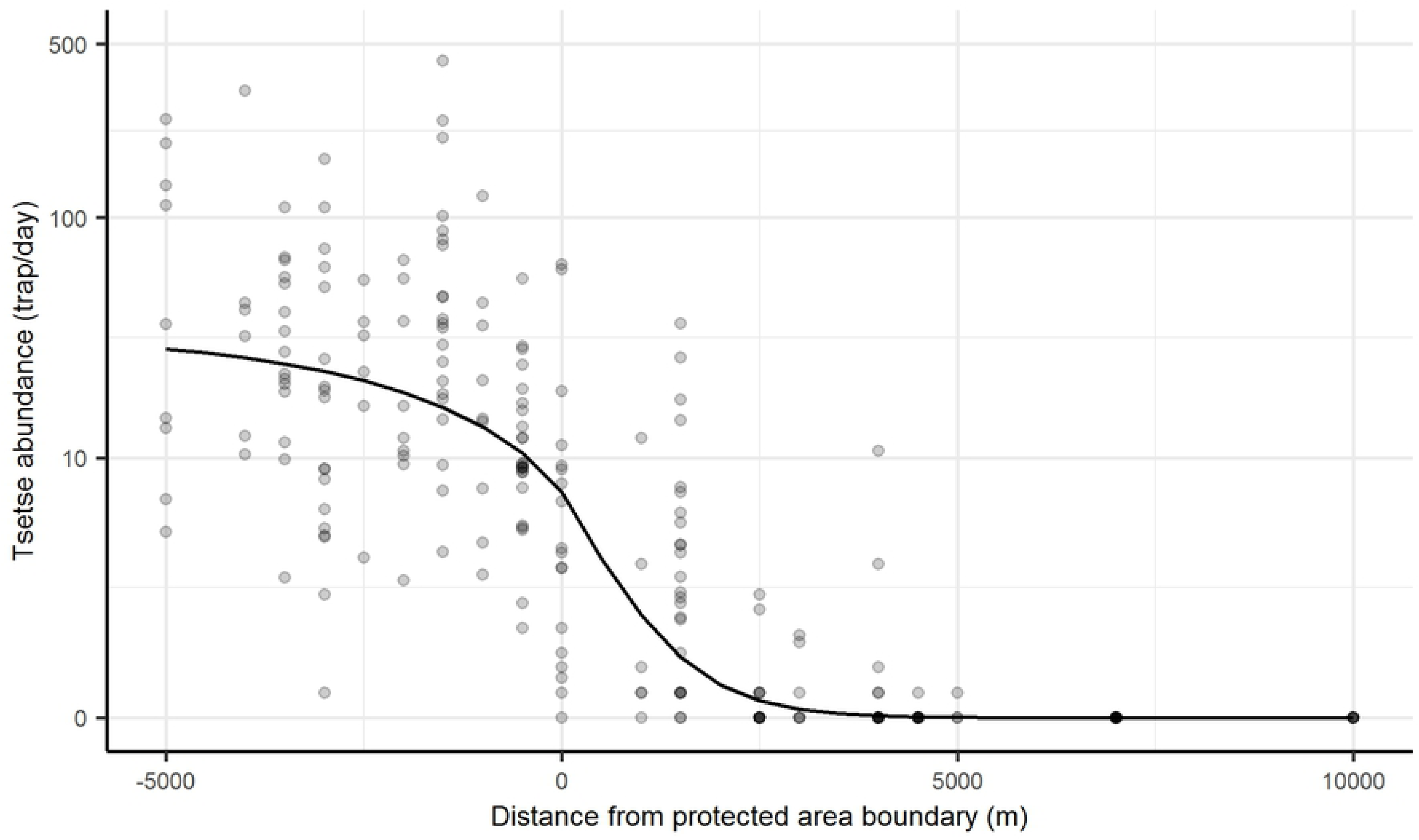
Modelled decline in tsetse abundance across the wildlife-livestock interface. Model fitted by nonlinear least squares regression to mean daily tsetse caught per trap across three surveys in 2015. Negative distances on *x* axis indicate inside wildlife areas where no additional mortality was modelled. The *y* axis is on log scale. Darker points indicate samples from multiple traps at the same distance.

### Simulating trypanosome transmission across the wildlife-livestock interface

Of the parameters detailed in Table 2, host incubation, host probability of infection and probability of recovery had the biggest effect on prevalence of trypanosomes in hosts, while the proportion of infected hosts that recover per day, and host-to-vector transmission probabilities had the biggest effect on prevalence of trypanosomes in vectors (S4 Fig, S5 Fig). From sensitivity analysis, of 1000 simulations with different parameter values, 138 had tsetse prevalence within the confidence intervals of that observed for *T. brucei s.l.* and 150 for *T. congolense*. Using these remaining parameter combinations, with the estimated additional mortality, *T. brucei* prevalence in hosts was on average 9.8% at 1 km from wildlife areas across simulations, declining to an average 4.0% by 2 km outside of wildlife areas across simulations, but *T. congolense* prevalence was on average 45.1% at 1 km outside of wildlife areas and 27.7% by 2 km across simulations (Fig 3).

**Fig 3.**
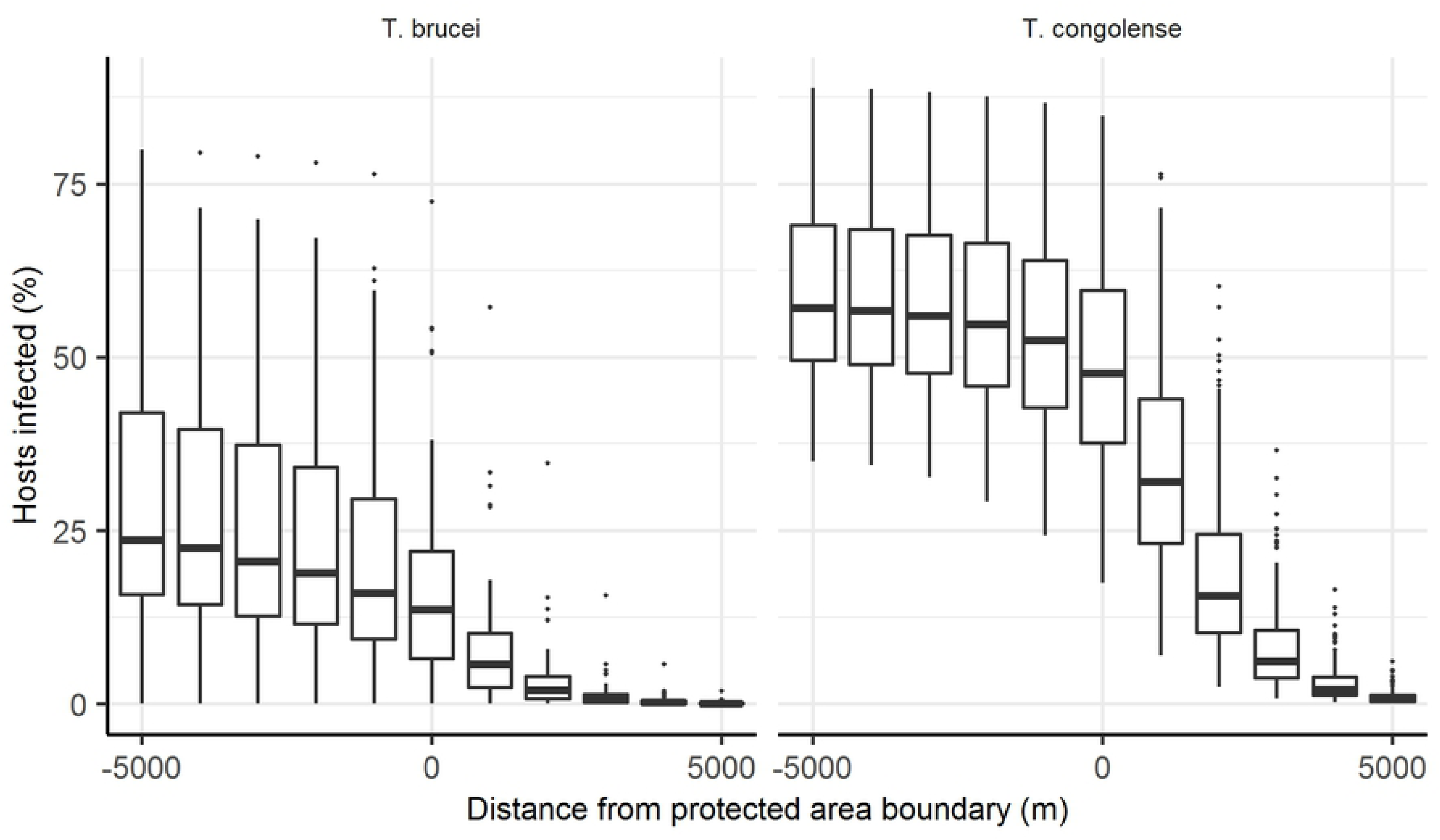
Modelled decline in trypanosome prevalence across the wildlife-livestock interface. Assuming additional probability of tsetse mortality/day in farming areas to be 0.152 as per model fits to the observed tsetse data, assuming tsetse disperse on average 500 m/day. The solid horizontal line in each boxplot shows the mean output from model runs using combinations of parameter values from sensitivity analysis that could explain the observed tsetse prevalence and hinges represent 25th and 75th percentiles.

## Discussion

We report the use of pyrethroid-based insecticides by farmers in Serengeti District at a frequency sufficient to impact tsetse populations. Our results support the findings of Ngumbi et al. [16] who reported the use of pyrethroids by farmers in Pangani, Myomero and Korogwe districts of Tanzania. To our knowledge, however, our study is the first to report farmer-led tsetse control, co-incident with tsetse decline and relatively low prevalence of *T. brucei s.l.* in cattle. There are other examples of insecticide-treated cattle being used to control tsetse and trypanosomiasis, but these were implemented by commercial ranches or with strong support from government institutions or donors [17–20]. Further detail on the scale of use across Tanzania, and why individual farmers are choosing to treat their cattle warrant further investigation.

Coupling questionnaires with hair sample analysis as we did in this study would be beneficial in further investigations. Questionnaires may be useful for gathering information on use, but issues with product labelling, including language translation, could result in inadequate application [21]. This may explain differences between reported insecticide use and quantified amounts on hair. The use of gas chromatography-mass spectrometry for analysis of livestock hair samples is expensive and a more cost-effective method for quantifying insecticide concentrations would be beneficial for future studies to aid larger-scale assessments of actual use.

With respect to the increased tsetse mortality in farming areas, due to uncertainty in both the data and model estimates, it was not possible to separate out mortality due to either insecticide-treated cattle or habitat degradation. A better understanding of the relative contribution of habitat degradation to tsetse decline at wildlife-livestock interface areas would help to identify where and when insecticide-treated cattle would be most effective.

*T. brucei s.l.* and *T. b. rhodesiense* prevalence, observed in cattle in Serengeti District during 2001, suggested to Kaare et al. [2] that r-HAT was re-emerging in this area. The *T. brucei s.l.* prevalence in our study was 1.25% (0.09 - 1.57) compared with 5.6% (3.78 – 7.94) estimated by Kaare et al. [2] and therefore there does not appear to have been an increase in risk in this area over time. Our modelling suggests that in areas of relatively high cattle density, such as our study site, where the majority of tsetse blood meals are from cattle, modest use of insecticide-treated cattle by livestock farmers can reduce the role of cattle in *T. b. rhodesiense* transmission despite the presence of high tsetse densities in adjacent wildlife areas. Treating cattle with pyrethroids may however be less effective against AAT [4]. Farmers at the boundary of wildlife areas are still therefore likely to treat their animals with trypanocides.

Our modelling involved several assumptions. We assumed that there was no overall change in tsetse population and trypanosome prevalence in wildlife areas over time. We did not account for seasonal changes in wild host movement which may influence prevalence in adjacent wildlife areas and therefore risk of infection in cattle. Nor did we account for trypanocide use, heterogeneity in insecticide-treated cattle use, or habitat quality in farming areas. These are likely important factors driving trypanosome prevalence. Our study does, however, extend the modelling carried out by Hargrove et al. [4] in being spatially-explicit and considering an interface context.

Treatment of cattle with insecticide offers the most cost-effective method of tsetse control [22] and in East Africa the risk of both tick- and tsetse-borne diseases of livestock provides a strong incentive for livestock keepers to treat their cattle regularly [23]. Effective control of savanna tsetse requires interventions conducted over large (>100 km^2^) areas [24]. This is possible for large commercial ranches [17,18] but much more difficult to implement and sustain with small-scale livestock farmers without co-ordination and financial support from donor or government agencies. Our findings, however, provide evidence that small-scale farmers can be enabled to control r-HAT. It is important to understand why farmers in Serengeti have adopted this strategy. For example, if ticks and tick-borne diseases are a major driver, then sustainable options that mitigate against resistance in the tick vector would be a priority. Understanding the underlying social, economic and political drivers of this phenomenon may lead us to the elusive goal of sustainable and cost-effective control of trypanosomiasis in east and southern Africa.

## Acknowledgements

The authors thank the field staff at Vector and Vector-Borne Diseases Research Institute, Tanzania. Permits were acquired from COSTECH and TAWIRI, Tanzania. The authors would also like to thank Louise Matthews and Shaun Keegan for providing review of the manuscript.

## Supporting Information Captions

**S1 Fig. Study site location.**

**S2 Fig. Scatter plots showing the relationship between model parameters and output.**

**S3 Fig. Partial rank correlation coefficient for each parameter in the tsetse population dynamics model.**

**S4 Fig. Results of sensitivity analysis for the trypanosome transmission model.**

**S5 Fig. Partial rank correlation coefficients for the trypanosome transmission model.**

**S1 Text. Additional methods.**

**S2 Text. Model equations.**

**S3 Text. Model sensitivity analysis and model fitting.**

**S1 Table. Fitted model parameter values**

## References

1. Auty H, Morrison LJ, Torr SJ, Lord J. Transmission dynamics of Rhodesian sleeping sickness at the interface of wildlife and livestock areas. Trends Parasitol. 2016;32: 608–621. doi:10.1016/j.pt.2016.05.003

2. Kaare MT, Picozzi K, Mlengeya T, Fèvre EM, Mellau LS, Mtambo MM, et al. Sleeping sickness - a re-emerging disease in the Serengeti? Travel Med Infect Dis. 2007;5: 117–24. doi:10.1016/j.tmaid.2006.01.014

3. Simarro PP, Cecchi G, Franco JR, Paone M, Diarra A, Ruiz-Postigo JA, et al. Estimating and mapping the population at risk of sleeping sickness. PLoS Negl Trop Dis. 2012;6: e1859. doi:10.1371/journal.pntd.0001859

4. Hargrove JW, Ouifki R, Kajunguri D, Vale G a, Torr SJ. Modeling the control of trypanosomiasis using trypanocides or insecticide-treated livestock. PLoS Negl Trop Dis. 2012;6: e1615. doi:10.1371/journal.pntd.0001615

5. Lord JS, Torr SJ, Auty HK, Brock PM, Byamungu M, Hargrove JW, et al. Geostatistical models using remotely-sensed data predict savanna tsetse decline across the interface between protected and unprotected areas in Serengeti, Tanzania. J Appl Ecol. 2018;55: 1997–2007. doi:10.1111/1365-2664.13091

6. Torr SJ, Maudlin I, Vale GA. Less is more: Restricted application of insecticide to cattle to improve the cost and efficacy of tsetse control. Med Vet Entomol. 2007;21: 53–64. doi:10.1111/j.1365-2915.2006.00657.x

7. Robinson TP, William Wint GR, Conchedda G, Van Boeckel TP, Ercoli V, Palamara E, et al. Mapping the global distribution of livestock. PLoS One. 2014;9. doi:10.1371/journal.pone.0096084

8. Davey BYJB. The outbreak of human trypanosomiasis (Trypanosoma rhodesiense infection) in Mwanza district, Tanganyika territory. Trans R Soc Trop Med Hyg. 1924;17: 474–481.

9. Jelinek T, Bisoffi Z, Bonazzi L, van Thiel P, Bronner U, de Frey A, et al. Cluster of African trypanosomiasis in travelers to Tanzanian national parks. Emerg Infect Dis. 2002;8: 634–5. doi:10.3201/eid0806.010432

10. Ripamonti D, Massari M, Arici C, Gabbi E, Farina C, Brini M, et al. African sleeping sickness in tourists returning from Tanzania: the first 2 Italian cases from a small outbreak among European travelers. CID. 2002;34: e19–22.

11. Lea R. Ecology and control of tsetse at the interface of conservation and farming areas in northern Tanzania. University of Liverpool. 2019.

12. Hargrove J, Williams B. A cost-benefit analysis of feeding in female tsetse. Med Vet Entomol. 1995;9: 109–119.

13. Muturi CN, Ouma JO, Malele II, Ngure RM, Rutto JJ, Mithöfer KM, et al. Tracking the feeding patterns of tsetse flies (Glossina genus) by analysis of bloodmeals using mitochondrial cytochromes genes. PLoS One. 2011;6: e17284. doi:10.1371/journal.pone.0017284

14. Rock KS, Torr SJ, Lumbala C, Keeling MJ. Quantitative evaluation of the strategy to eliminate human African trypanosomiasis in the Democratic Republic of Congo. Parasit Vectors. 2015; 1–13. doi:10.1186/s13071-015-1131-8

15. Dale C, Welburn SC, Maudlin I, Milligan PJM. The kinetics of maturation of trypanosome infections in tsetse. Parasitology. 1995;111: 187. doi:10.1017/S0031182000064933

16. Ngumbi AF, Silayo RS. A cross-sectional study on the use and misuse of trypanocides in selected pastoral and agropastoral areas of eastern and northeastern Tanzania. Parasites and Vectors. Parasites & Vectors; 2017;10: 1–9. doi:10.1186/s13071-017-2544-3

17. Fox RGR, Mmbando SO, Fox MS, Wilson A. Effect on herd health and productivity of controlling tsetse and trypanosomosis by applying deltamethrin to cattle. Trop Anim Health Prod. 1993;25: 203–214. doi:10.1007/BF02250869

18. Baylis M, Stevenson P. Trypanosomiasis and tsetse control with insecticidal pour-ons - Fact and fiction? Parasitol Today. 1998;14: 77–82. doi:10.1016/S0169-4758(97)01170-8

19. Warnes ML, Van Den Bossche P, Chihiya J, Mudenge D, Robinson TP, Shereni W, et al. Evaluation of insecticide-treated cattle as a barrier to re-invasion of tsetse to cleared areas in northeastern Zimbabwe. Med Vet Entomol. 1999;13: 177–184. doi:10.1046/j.1365-2915.1999.00148.x

20. Hargrove J, Omolo S, Msalilwa J, Fox B. Insecticide-treated cattle for tsetse control: The power and the problems. Med Vet Entomol. 2000;14: 123–130. doi:10.1046/j.1365-2915.2000.00226.x

21. Allan F. East Coast fever and vaccination at the livestock/wildlife interface. University of Edinburgh. 2018.

22. Shaw APM, Torr SJ, Waiswa C, Cecchi G, Wint GRW, Mattioli RC, et al. Estimating the costs of tsetse control options: An example for Uganda. Prev Vet Med. Elsevier B.V.; 2013;110: 290–303. doi:10.1016/j.prevetmed.2012.12.014

23. Muhanguzi D, Picozzi K, Hatendorf J, Thrusfield M, Welburn SC, Kabasa JD, et al. Collateral benefits of restricted insecticide application for control of African trypanosomiasis on Theileria parva in cattle: A randomized controlled trial. Parasites and Vectors. 2014;7: 1–10. doi:10.1186/1756-3305-7-432

24. Torr SJ, Hargrove JW, Vale G a. Towards a rational policy for dealing with tsetse. Trends Parasitol. 2005;21: 537–41. doi:10.1016/j.pt.2005.08.021

